# Preserved neural population dynamics across animals performing similar behaviour

**DOI:** 10.1101/2022.09.26.509498

**Authors:** Mostafa Safaie, Joanna C. Chang, Junchol Park, Lee E. Miller, Joshua T. Dudman, Matthew G. Perich, Juan A. Gallego

## Abstract

Animals of the same species often exhibit similar behaviours that are advantageously adapted to their body and their environment. These behaviours are shaped by selection pressures over evolutionary timescales at the species level, yet each individual produces these behaviours using a different, uniquely constructed brain. It remains unclear how these common behavioural adaptations emerge from the idiosyncratic neural circuitry of a given individual. Here, we hypothesised that the adaptive behaviour of a species requires specific neural population ‘latent dynamics’. These latent dynamics should thus be preserved and identifiable across individuals within a species, regardless of the idiosyncratic aspects of each individual’s brain. Using recordings of neural populations from monkey and mouse motor cortex, we show that individuals from the same species share surprisingly similar neural dynamics when they perform the same behaviour. The similarity in neural population dynamics extends beyond cortical regions to the dorsal striatum, an evolutionarily older structure, and also holds when animals consciously plan future movements without overt behaviour. These preserved dynamics are behaviourally-relevant, allowing decoding of intended and ongoing movements across individuals. We posit that these emergent neural population dynamics result from evolutionarily-imposed constraints on brain development, and reflect a fundamental property of the neural basis of behaviour.

## Introduction

Animals of the same species are capable of learning and executing a certain repertoire of adaptive behaviours. For example, all macaque monkeys move their arms in similar ways when grabbing fruit from a branch or when reaching for a target on a computer screen in a laboratory. Ultimately, these behaviours are driven by the coordinated activity of neural populations throughout the brain. This activity emerges from the ‘latent dynamics’, which are the time-dependent activation of the dominant patterns of neural co-variation^1–3^. These latent dynamics seem to be shaped by circuit and biophysical constraints^4–7^. Given the large differences in brain circuitry across individuals from the same species —including in number of type-specific neurons, dendritic morphology, and receptor distribution^8–12—^ it remains unclear how similar adaptive behaviours emerge from such idiosyncratic neural circuitry. One possibility is that unique circuits in each individual generate unique neural dynamics to produce the same behaviour. Indeed, the high degrees of freedom of neural activity relative to behaviour^13, 14^ allow distinct latent dynamics to produce similar behavioural output. Alternatively, the behaviour could be tied to specific latent dynamics. In this case, the idiosyncratic neural circuits must be tuned to produce the necessary emergent latent dynamics needed to generate the behaviour.

We hypothesised that preserved neural population latent dynamics are the basis of species-specific behaviours: different individuals from the same species engaged in the same behaviour generate latent dynamics that each is a different instantiation of a common trajectory through a species-wide neural landscape (Figure 1). Our hypothesis provides several testable predictions. First, since low-level details of neural circuits are idiosyncratic, they should not be necessary to account for the emergence of species-typical behaviours. Accordingly, different animals of the same species engaged in the same behaviour should exhibit preserved latent dynamics. Second, the extent of preservation of the latent dynamics across individuals should be constrained by the similarity of the behavioural output. Third, since low-dimensional latent dynamics are found throughout the brain, not just in cortical regions^15–17^, we also expect structures such as the basal ganglia, which have co-evolved with cortex for hundreds of millions of years^18^, to exhibit shared latent dynamics across animals performing the same behaviour. Fourth, since covert behaviours seem to be mediated by the same neural circuits as overt behaviours^19–21^, we should find shared latent dynamics across animals performing the same cognitive task.

**Figure 1:**
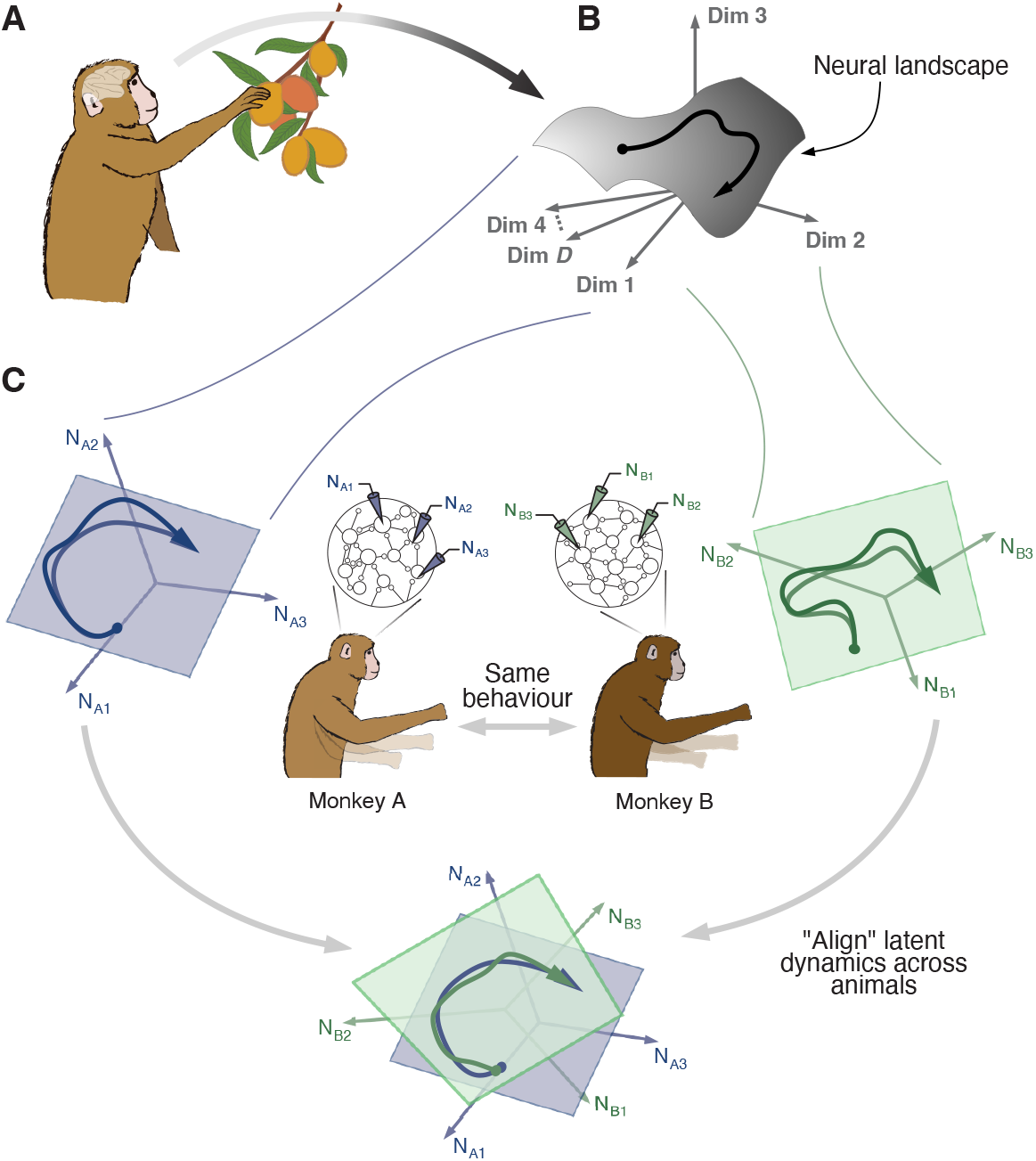
Hypothesis. Different individuals from the same species performing the same behaviour (Panel **A**) will generate preserved neural population latent dynamics by instantiating a species-wide ‘neural landscape’ (Panel **B**). These preserved latent dynamics can be revealed by ‘aligning’ the individual-specific latent dynamics estimated from their neural population recordings (Panel **C**).

## Results

We tested the four predictions outlined above using neural population recordings from monkeys and mice as they performed upper limb tasks. First, we analysed motor cortical recordings from monkeys engaged in an instructed-delay centre-out reaching task with eight targets (Figure 2A, Figure S1A, Methods). All three monkeys were well-trained in the task and exhibited highly stereotyped hand trajectories (mean trajectory correlation for each monkey: *r* = 0.89, 0.90, and 0.92, Figure S1B). For each session, we used Principal Component Analysis (PCA) to estimate the latent dynamics underlying overt movement execution by projecting the firing rates of each recorded neuron (or multi-unit, for monkey *J*) onto the leading ten PCA axes (the ‘neural modes’; examples in Figure S1C; Methods). We then ‘aligned’ the latent dynamics of each pair of experimental sessions from two different animals using Canonical Correlation Analysis (CCA^22^), a method that finds linear transformations that maximise the correlations between two sets of signals (similar to Ref.^23^–^25^; see Figure S1D).

**Figure 2:**
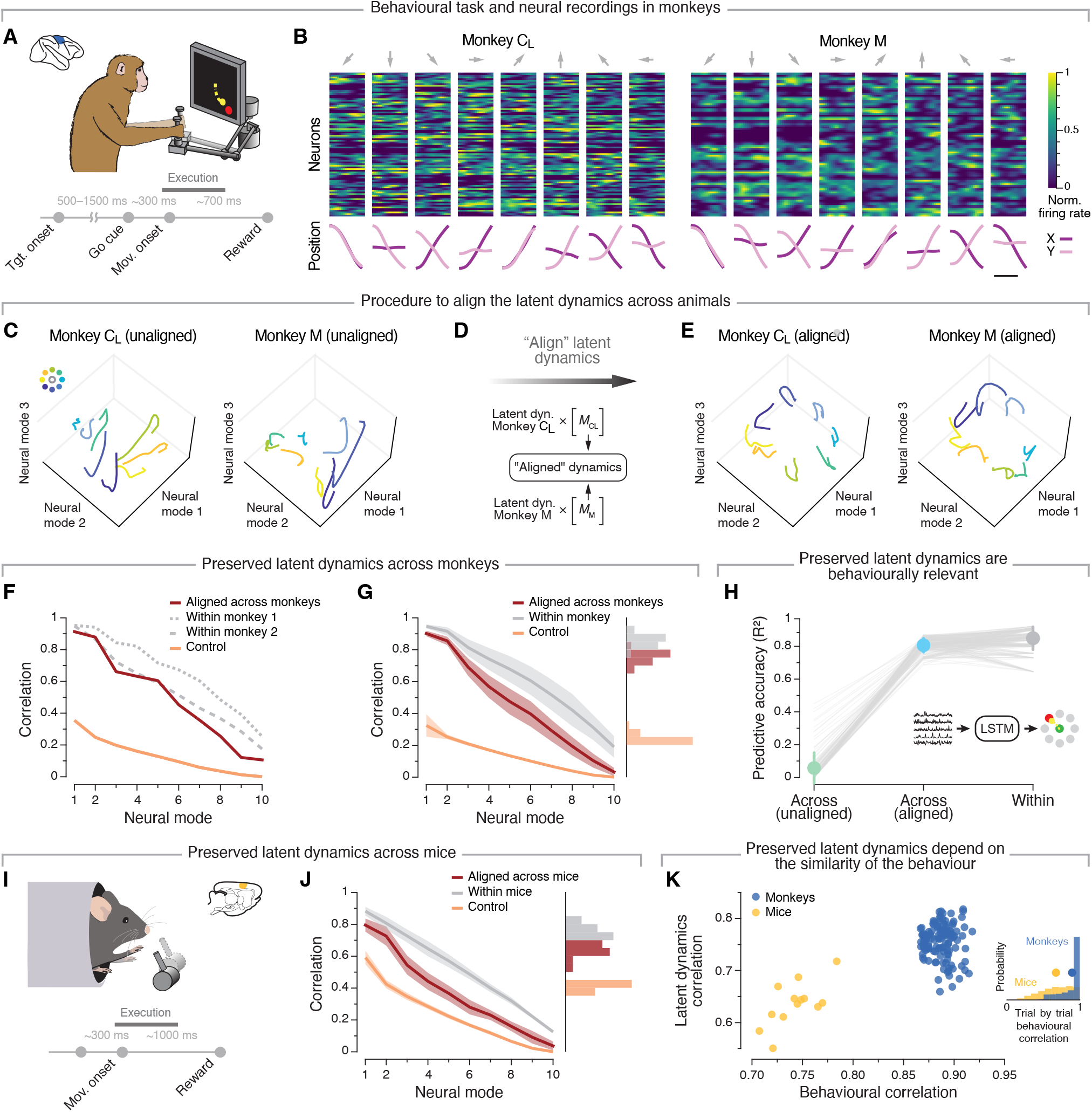
Preservation of latent dynamics across animals performing the same behaviour. **A)** Monkeys performed an eight-target instructed-delay centre-out reaching task using a planar manipulandum. **B)** Example normalised neural firing rates aligned to movement onset for two different monkeys (top) and corresponding hand trajectories (bottom). Each column, one reach to each of the eight targets. Arrows on top of each column, direction of the target. Scale bar, 300ms. **C)** Three-dimensional representation of the motor cortical latent dynamics for each of the two monkeys plotted in Panel B before alignment using CCA (Panel **D**). **E)** Alignment using CCA revealed strong similarities in the latent dynamics across monkeys. Same data as in Panel B. Axes are orthogonolised for presentation only (Methods). **F)** Correlations of the aligned (red) latent dynamics for the example comparison shown in Panels C–E. Note that the correlation between aligned latent dynamics across monkeys was quite close to the within-monkey correlations (grey), and largely exceeded our lower-bound control (orange). **G)** Similarity in latent dynamics across all pairs of sessions from three different monkeys. Data include 21 sessions and 126 pairs of comparisons. Histograms on the right show the distribution of the mean correlation across the leading four dimensions. Line and shaded area, mean±s.d. **H)** Decoding of hand position based on the latent dynamics. Decoders trained on the aligned latent dynamics from one monkey allowed predicting hand position from a different monkey (blue) virtually as well as decoders trained and tested within the same session (grey), while decoding was poor without alignment. Data points, individual comparisons between two sessions from different monkeys (*n* = 126). Errorbars, mean*±*s.d. **I)** Mice grasped and pulled a joystick located in one of two positions (left or right) to receive a reward. **J)** Similarity in latent dynamics across mice performing the grasping and pulling task. Data include six sessions across four different mice, formatted as in Panel G. **K)** The similarity in latent dynamics across animals is related to the similarity of their behaviour. The plot shows, for each pair of sessions, the mean of the top four canonical correlations between the latent dynamics as a function of the mean correlation of the behaviour. Single dots, pairs of sessions colour coded by species. Inset: Distribution of the behavioural correlations for all pairs of trials from every two different mice and monkeys. Circles, mean.

This linear method revealed that the ostensibly different latent dynamics of two different monkeys are indeed highly preserved (Figure 2E): across-animal correlations greatly exceeded a lower bound established by comparing two randomly selected behavioural epochs (‘Control’ in Figure 2F). Most importantly, these correlations were nearly as high as values obtained by aligning two subsets of trials within a single session from the same animal (‘Within’ in Figure 2F; additional examples in Figure S2A). This result held across all pairs of sessions from all three monkeys (*n* = 126 sessions, Figure 2G), and did not depend on the assumed dimensionality of the neural manifold (Figure S3).

While we have demonstrated the conservation of latent dynamics across animals, these shared dynamics may not necessarily be relevant to behaviour. To address this, we trained neural network decoders (Long Short-Term Memory networks, or LSTMs) to predict the hand trajectories of one animal^26^ and tested their performance on a second animal (Methods). The across-animal decoding accuracy approached the upper bound provided by the performance of decoders trained and tested on the same animal (Figure 2H; example predictions in Figure S4). Thus, the shared motor cortical dynamics across animals do contain detailed information about the ongoing movement kinematics.

We then analysed data from four mice trained to perform a grasping and pulling joystick task (Figure 2I and Figure S5). We found that both within and across individuals, the behavioural output was less similar from trial to trial than for the monkey dataset (compare Figure S1A,B to Figure S5A,B). As predicted, our alignment procedure revealed that motor cortical latent dynamics were largely preserved across mice (Figure 2J and Figure S2B; example in Figure S6A). Yet, the extent of these correlations was lower than those of monkeys’ (compare Figure 2G to Figure 2J), which directly impacted our ability to decode movement kinematics across animals (Figure S7). We predicted that much of this difference could be explained by the more highly stereotyped behaviour of the monkeys compared to the mice (Figure 2K-inset). Comparing between species confirmed that behavioural stereotypy was indeed well associated with both the preservation of the latent dynamics across individuals (Figure 2K) and across-animal decoding accuracy (Figure S8). These results demonstrate in two evolutionarily divergent species that there is a direct correspondence between the similarity of behavioural output and the preservation of motor cortical latent dynamics across individuals.

We performed several control analyses to confirm that this close relationship between behavioural stereotypy and preservation of latent dynamics across individuals from the same species is not a trivial consequence of our methodology. First, we compared our results with a previous study that investigated the dynamics of motor cortical activity within an individual across different but related wrist manipulation and reach-to-grasp tasks^27^. The latent dynamics of different monkeys performing the same task are much more preserved than those of the same monkey performing two distinct but related hand motor tasks that engage the same cortical circuits (Figure S9). Moreover, as we have previously shown, preserved latent dynamics cannot be explained by stable movement tuning, nor do they persist following nonlinear transformations^24^. These findings show that the alignment method alone, even within individuals, does not suffice to generate preserved latent dynamics. Lastly, our hypothesis requires that behavioural similarity is necessary but not sufficient to allow for alignment of neural dynamics. In particular, we would not expect brains with different circuit properties to share very similar latent dynamics even if the behaviour is the same. To test this, we trained recurrent neural networks to perform the centre-out reaching task to study the relationship between latent dynamics and behavioural output. By modifying how a subset of networks were trained (Methods), we could create pairs of models that generated highly similar behaviour but exhibited latent dynamics that were distinct even after alignment (Figure S10). Thus, preservation of latent dynamics is not just a trivial consequence of behavioural similarity; instead, it also likely reflects fundamental constraints in the underlying circuit implementation that are shared by different brains of different individuals from the same species.

Given that the motor cortex is the main cortical output to the spinal circuits that generate movement, the close correspondence between motor cortical latent dynamics and behavioural output may uniquely result from the architecture and projections of this region. To test whether preserved latent dynamics exist across the brain, we studied the subcortical nuclei of basal ganglia, which do not directly project to spinal cord but are crucial for various aspects of behaviour^28–34^. We predicted that basal ganglia latent dynamics would also be preserved across animals performing the same task. Replicating our alignment analysis on neural population recordings from mouse dorsolateral striatum during a grasping and pulling task (Figure 3A) revealed preserved latent dynamics across individuals (Figure 3B and Figure S2C; example in Figure S6B). Moreover, despite the vast differences in circuit and cellular architecture between cortex and striatum^34, 35^, both the across-animal correlations (compare Figure 3B and Figure 2J) and the across-animal decoding performance of hand trajectories (Figure S8) were equally large for both regions. Therefore, stable latent dynamics across animals performing the same behaviour are not confined to motor cortical regions—they likely extend to different regions throughout the brain, including an evolutionarily older structure that is shared among all vertebrates^36^.

**Figure 3:**
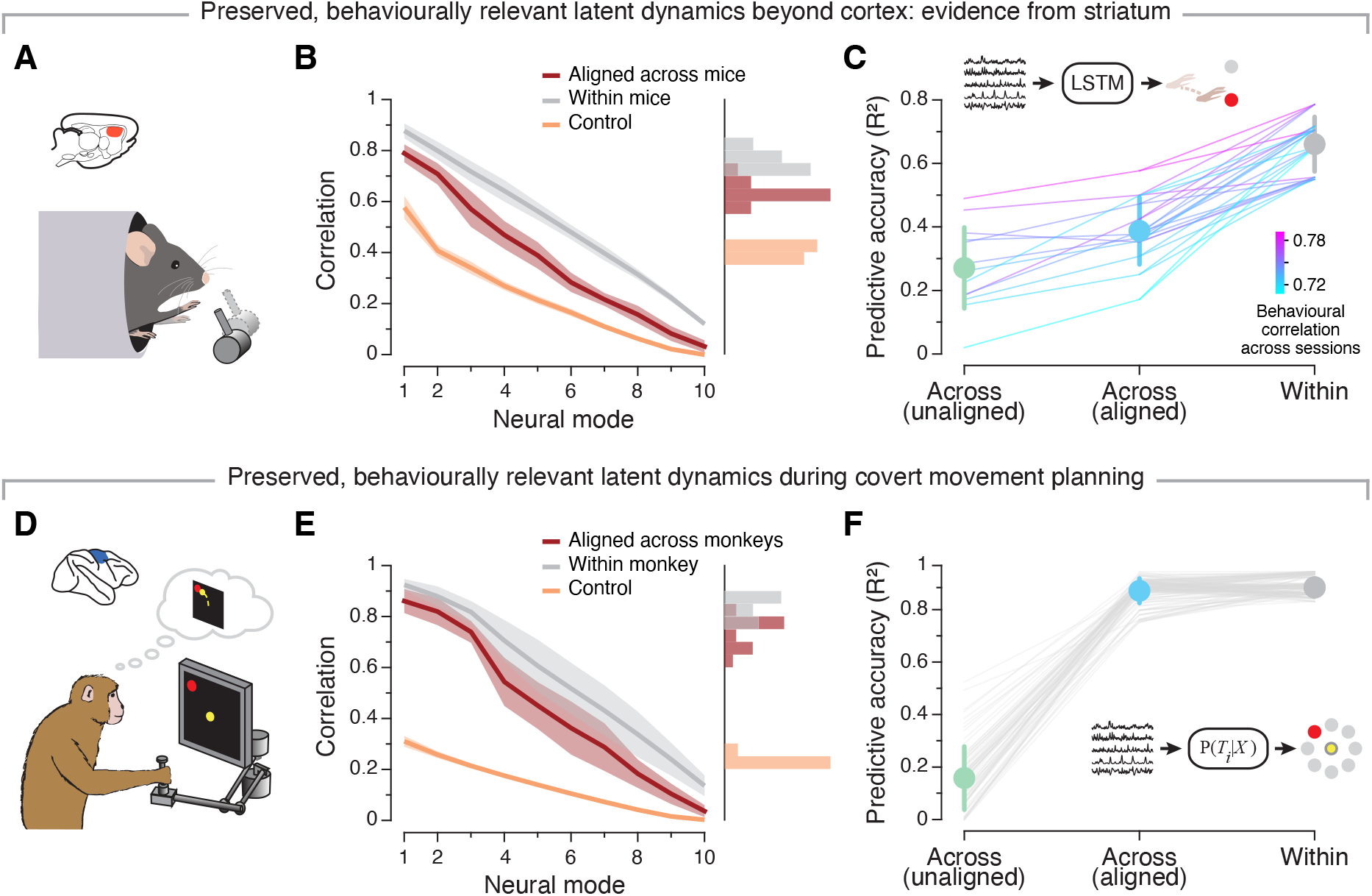
Preserved latent dynamics extend beyond cortical regions to the evolutionarily older basal ganglia, and are also present during covert behaviour. **A)** We studied whether shared latent dynamics can also be found in subcortical regions by analysing recordings from mouse dorsolateral striatum during a grasping and pulling task. **B)** Similarity in striatal latent dynamics across mice performing the same grasping and pulling task. Data include six sessions across four mice and are presented as in Figure 2G. **C)** Hand trajectories can be decoded across mice using their shared striatal latent dynamics. Data presented as in Figure 2H. **D)** To investigate the preservation of latent dynamics during covert behaviour, we examined the preparatory period preceding movement execution in the monkey reaching task. **E)** Similarity in motor cortical latent dynamics when preparing to reach to a target. Data include 18 sessions and *n* = 72 pairs of comparisons across two different monkeys, and are presented as in Figure 2G. **F)** The preserved latent dynamics during covert behaviour contain behaviourally-relevant information. Näive Bayes classifiers trained on the aligned latent dynamics (blue) predicted the intended target in a different monkey virtually as well as classifiers trained and tested on the same monkey (grey), while classification was poor without alignment (green).

We have shown the preservation of latent dynamics across brain regions during active, overt behaviour. However, higher-order animals also engage in a variety of covert behaviours such as deliberation and planning. These processes require neural activity that is predominantly generated internally by the brain. Given that such covert behaviours involve brain regions that also mediate overt behaviours^19^–21, we predicted that the latent dynamics underlying these more cognitive processes would also be shared across individuals of the same species. We tested this prediction by analysing motor cortical activity as monkeys planned an upcoming movement prior to executing it (Figure 3D). The latent dynamics during the instructed delay period were highly correlated across animals (Figure 3E and Figure S2D; example in Figure S6C), and were virtually identical to those during overt reaching behaviour (*cf*. Figure 2). Moreover, these aligned latent dynamics were also predictive of behaviour: Bayesian models predicting the upcoming reaching target based on the aligned latent dynamics from one monkey generalised to another (Figure 3F; Methods). Thus, different individuals use preserved latent dynamics not only to execute the same movement but also to perform the same covert mental process. This result also strengthens our previous observation of preserved latent dynamics during overt behaviour: afferent feedback arriving at the motor cortex^37^, 38 may contribute to the observed similarity in latent dynamics during overt movement, yet the latent dynamics are entirely internally generated during covert processes such as movement planning.

## Discussion

Neural population latent dynamics have been proposed as first-level explainers of behavioural and cognitive functions^1–3^, a view that has shed light onto the neural basis of numerous phenomena, such as processes underlying covert^39, 40^ and overt behaviour^27, 41^, how learning may happen in neural circuits^4, 42^, and how information may flow between different brain regions^38, 43, 44^. Here, we extend recent works^13, 39, 45–47^ to show that latent dynamics are shared across different individuals engaged in the same behaviour, providing a novel framework to study brain function across populations of animals. Our results are important for the development of brain-controlled devices such as neuroprosthetics^48, 49^: with proper alignment, decoders trained on one participant could be readily translated to other individuals to restore movement^50–54^ or even cognitive^55^ or affective^56^ functions.

Finally, our findings also have broad implications for the study of the neural basis of behaviour. We introduced two competing possibilities to explain how similar behaviour can emerge from different neural circuitry. In the first, the unique circuitry of each individual could produce individual-specific latent dynamics. This would imply that the appropriate mapping from these idiosyncratic latent dynamics must be learned during development of the individual in order to enable the desired behaviour. In the second hypothesis, the idiosyncratic circuitry of each individual still produces preserved latent dynamics tied to a specific behaviour. In this case, the appropriate mapping from circuit to latent dynamics would be an emergent property that is genetically specified. Our results support this second hypothesis, and raise an intriguing possibility: given that in higher vertebrates the genome does not specify the implemented architecture in great detail, e.g., to the level of the synapse^57^–59, the genome may provide a ‘generative model’ that is instantiated by each individual’s brain. This generative model may constrain low-level details of the neural circuitry such that the neural population latent dynamics required for the behavioural repertoire emerge throughout development.

## Methods

### Subjects and the behavioural tasks

#### Monkeys

We trained three monkeys (monkeys *C, M*, and *J*; all male, *Macaca mulatta*) to sit in a primate chair and make reaching movements using a customised planar manipulandum. All three monkeys were trained to perform a similar two-dimensional centre–out reaching task for several months prior to the neural recordings. In the task, the monkey moved its hand to the centre of the workspace to begin each trial. After a variable waiting period, the monkey was presented with one of eight outer targets, equally spaced in a circle and selected randomly with uniform probability. Then, an auditory go cue signalled the animals to reach to the target. Monkeys were required to reach the target within 1 s after the go cue and hold for 0.5 s to receive a liquid reward, except for monkey *J*, who was trained without the 0.5 s target hold time and therefore made larger movements (Figure S1A, right). As the monkeys performed this task, we recorded the position of the endpoint at a sampling frequency of 1 kHz using encoders in the joints, and digitally logged the timing of task events, such as the go cue. Portions of these data have been previously published and analysed in Refs.24, 25, 42, 60.

#### Mice

Four 8–16 week old mice, were trained to perform a forelimb grasping and pulling task (similar to ^34^, 61) for approximately one month, following habituation to head-fixation and the recording setup. In each trial, mice had to reach and pull a joystick positioned approximately 1.5 cm away from the initial hand position. The joystick appeared, without any cue, in one of two positions (left or right, < 1 cm apart). Mice could then self-initiate a reach to the joystick and pull it inwards to get a liquid reward. The joystick was weighted with either a 3 g or a 6 g load (light or heavy), making up a total of 4 trial types (2 joystick positions × 2 loads). Each trial type was repeated 20 times before task parameters were switched to the next trial type without any cue. Each session consisted of two repetitions of each set of four trial types presented in the same order, making up 2 × 4 × 20 = 160 trials. In this work, light and heavy trials were pooled together, since every analysis is aligned on the onset of the reaching phase and the location of the joystick does not depend on its weight.

### Neural recordings

#### Monkeys

All surgical and experimental procedures were approved by the Institutional Animal Care and Use Committee of Northwestern University under protocol #IS00000367. We implanted 96-channel Utah electrode arrays in the primary motor cortex (M1) or dorsal premotor cortex (PMd) using standard surgical procedures. Throughout the paper, neural recordings from these two subregions of the motor cortex were pooled together and denoted as motor cortex. This allowed us to ensure we could evaluate overt and covert dynamics within the same population. Implants were done in the opposite hemisphere of the hand animals used in the task. Monkey *C* received two sets of implants: a single array in the right M1 while performing the task using the left hand, and later, arrays in the left M1 and PMd while using the right hand (respectively, monkeys *C*_*R*_ and *C*_*L*_ in our previous works^24^). Note that for all across-individual analyses, *C*_*R*_ and *C*_*L*_ are considered the same animal.

Neural activity was recorded during the behaviour using a Cerebus system (Blackrock Microsystems). The recordings on each channel were band-pass filtered (250 Hz–5 kHz) and then converted to spike times based on threshold crossings. The threshold was set to 5.5 × the root-mean square activity on each channel. We also manually spike sorted the recordings from monkeys *C* and *M* to identify putative single neurons. Monkey *J* had fewer well-isolated single units than the other monkeys, so rather than spike sorting we directly applied the multi-unit threshold crossings acquired on each electrode. However, it has been shown that the latent dynamics estimated from multi-unit and single neuron activity are similar^62^, an observation that holds true for aligning latent dynamics with CCA^24^ (note that we refer to both single neurons and multi-units simply as units). We included multiple experimental sessions from each monkey: 8 from *C*_*L*_, 4 from *C*_*R*_, 6 from *M*, and 3 from *J*. These experimental sessions were chosen based on the high number of units and blind to the behaviour of the animal. A more detailed description of the behavioural and neural recording methods is presented in^24^.

#### Mice

All surgical and experimental procedures were approved by the Institutional Animal Care and Use Committee of Janelia Research Campus. A brief (<2 h) surgery was first performed to implant a 3D-printed headplate^63^. Following recovery, the water consumption of the mice was restricted to 1.2 ml per day, in order to train them in the behavioural task. Following training, a small craniotomy for acute recording was made at 0.5 mm anterior and 1.7 mm lateral relative to bregma in the left hemisphere. A Neuropixels probe was centred above the craniotomy and lowered with a 10 degree angle from the axis perpendicular to the skull surface at a speed of 0.2 mm/min. The tip of the probe was located at 3 mm ventral from the pial surface. After a slow and smooth descent, the probe was allowed to sit still at the target depth for at least 5 min before initiation of recording to allow the electrodes to settle.

Neural activity was filtered (high-pass at 300 Hz), amplified (200*×* gain), multiplexed and digitised (30 kHz), and recorded using the SpikeGLX software (https://github.com/billkarsh/SpikeGLX). Recorded data were pre-processed using an open-source software KiloSort 2.0 (https://github.com/MouseLand/Kilosort) and manually curated using Phy (https://github.com/cortex-lab/phy) to identify putative single units in each of the primary motor cortex and dorsolateral striatum. A total of six experimental sessions (from four mice, see Figure S5) with simultaneous motor cortical and striatal recordings were included in this work.

### Data analysis

We used a similar approach for both monkey and mouse data. In all the analyses, we only considered the trials in which the animal successfully completed the task within the specified time and received a reward. In every analysis, an equal number of trials to each target was randomly selected (eight targets for the monkeys and two targets for mice). Trial order was randomised to eliminate the possible effect of the passage of time. Within each trial, we isolated a window of interest that captured most of the movement, starting 50 ms before movement onset and ending 400 ms after movement onset.

To analyse covert behaviour in monkeys, we used a window that spanned the movement planning period, which started 400 ms before movement onset and ended 50 ms after movement onset. To establish a ‘behaviourally irrelevant’ window as control, we randomly selected windows of similar length (450 ms) along the entire duration of the inter-trial and trial periods combined. Importantly, all of our results held when changing the analysis windows within a reasonable range. All the analyses were implemented in Python using open-source packages such as numpy, matplotlib, sci-kit, scipy, and pandas^64–68^ and custom code.

#### Behavioural correlation

To assess the behavioural stereotypy of a given animal, we calculated hand trajectory correlations (Pearson’s correlation) of every pair of trials within a session. The distributions in Figure 2I-inset illustrate these correlations pooled across all the sessions included in this work. To determine the behavioural similarity across pairs of sessions from different monkeys or mice (Figure 2I), we similarly calculated correlations to compare all pairs of trials from the two sessions.

#### Neural population latent dynamics

To estimate the latent dynamics associated with the recorded neural activity in each session for both mice and monkeys, we computed a smoothed firing rate as a function of time for each unit. We obtained these smoothed firing rates by applying a Gaussian kernel (*σ* = 50 ms) to the binned square-root transformed firings (bin size= 30 ms) of each unit. We excluded units with a low mean firing rate (< 1 Hz mean firing rate across all bins), but we did not perform any further exclusions, e.g., based on lack of modulation or behavioural tuning. For each session, this produced a neural data matrix *X* of dimension *n* by *T*, where *n* is the number of recorded units and *T* the total number of time points from all concatenated trials on a given day; *T* is thus given by the number of targets per day × number of trials per target × number of time points per trial. We performed this concatenation as described above after randomly sub-selecting the same number of trials for all targets for each animal (15 trials for monkeys, 22 for mice). For each session, the activity of *n* recorded units was represented as a neural space—an *n*-dimensional sampling of the space defined by the activity of all neurons in that brain region. In this space, the joint recorded activity at each time bin is represented as a single point, the coordinates of which are determined by the firing rate of the corresponding units. Within this space, we computed a low-dimensional neural manifold by applying PCA to *X*. This yielded *n* PCs, each a linear combination of the smoothed firing rates of all *n* recorded units. These PCs are ranked based on the amount of neural variance they explain. We defined an *m*-dimensional, session-specific manifold by only keeping the leading *m* PCs, which we referred to as neural modes. We chose a manifold dimensionality *m* = 10, based on previous studies examining motor cortical recordings during upper limb tasks^4, 24, 42^. Across all datasets, a ten-dimensional manifold explained *∼* 60% of the neural variance for each of the monkey motor cortex, mouse motor cortex, and mouse striatum (Figure S5). Our results held within a reasonable range of dimensionalities, similar to^24, 27, 42^ (see Figure S3). We computed the latent dynamics within the manifold by projecting the time-varying smoothed firing rates of the recorded neurons onto the *m* = 10 PCs that span the manifold. This produced a data matrix *L* of dimensions *m* by *T*.

#### Canonical correlation analysis

We addressed our hypothesis that different animals performing the same behaviour would share preserved latent dynamics by aligning the dynamics using Canonical Correlation Analysis (CCA)^22, 24^. CCA was applied to the latent dynamics of each pair of sessions after randomising the temporal order of trials and matching the number of trials to each target. For details on CCA, see^24^.

We measured the similarity in latent dynamics across animals by computing the across-animal correlations as the canonical correlations (CCs) across all pairs of sessions from any two different monkeys or mice. To establish how strong the across-animal correlations were, we computed an upper bound defined by the within-animal correlations, which we calculated as the 99th percentile of the correlations between two randomly selected subsets of trials within any given session, repeated 1000 times. The ‘control’ correlations represent a lower bound for the CCs and were computed similarly to the across-animal correlations, but with shuffling the targets across the two sessions and using a randomly selected control window (as described above) in each trial, instead of the movement or preparatory epoch.

Note that each point in Figure 2I denotes the mean of the top four canonical correlations of the latent dynamics and the mean trajectory correlation of all possible pairs of trials across two animals. Furthermore, when showing pairs of ‘aligned’ trajectories across animals, such as in the right hand side of Figure 2C and Figure S6, the CCA axes were orthogonalised using singular value decomposition for visualisation purposes.

#### Decoding analysis

To test whether the aligned latent dynamics maintain movement-related information, we built standard decoders to predict hand trajectory during overt behaviour. If the aligned latent dynamics across different animals were behaviourally relevant, they would allow predicting time-varying hand trajectories even if the methods used to identify them (PCA and CCA) are not supervised, that is, they do not attempt to optimise decoding performance. We compared the predictive accuracy of three different types of decoders: 1) a within-animal decoder trained and tested (using ten-fold cross-validation) on two non-overlapping subsets of trials from each session of each animal; 2) an across-animal ‘aligned’ decoder that was trained on the aligned dynamics from one animal and tested on another, a comparison we performed on each pair of sessions from two different animals; 3) an across-animal ‘unaligned’ decoder that was trained on the latent dynamics from one animal and tested on another without aligning the dynamics using CCA. We also performed a similar analysis to predict the upcoming target during covert movement preparation in monkeys (Figure 3C).

Hand trajectory decoders were LSTM models with two LSTM layers, each with 300 hidden units, followed by a linear output layer. The models were implemented with Pytorch^69^ and trained for 10 epochs with the Adam optimiser, with a learning rate of 0.001. Upcoming target classifiers were Gaussian Näive Bayes models (the GaussianNB class in^66^). We included three bins of recent latent dynamics history, for a total of 90 ms, in the input of both the decoders and the classifiers. These additional neural inputs incorporate information about intrinsic neural dynamics and account for transmission delays. The *R*^2^ value, defined as the squared correlation coefficient between actual and predicted hand trajectories, was used to quantify decoder performance. Hand trajectory was a two-dimensional signal in monkeys and a three-dimensional signal in mice. We built separate decoders to predict hand trajectories along the *X, Y* (and *Z* for mice) axes. We then reported the average performance across all axes. For target classification, we reported the mean accuracy of the classifier (the score() method).

### Recurrent neural network models

#### Model Architecture

To show that the preservation of latent dynamics across animals engaged in the same task is not a trivial byproduct of behaviour, we trained recurrent neural networks (RNNs) to perform the same centre-out reaching task as the monkeys. These models were implemented using Pytorch^69^. Similar to previous studies simulating motor cortical dynamics during reaching^70, 71^, we implemented the dynamical system 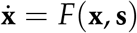 to describe the RNN dynamics:

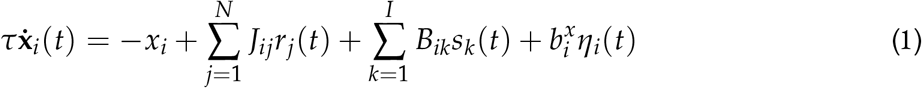

where *x*_*i*_ is the hidden state of the *i*th unit and *r*_*i*_a is the corresponding firing rate following *tanh* activation of *x*_*i*_. All networks had *N* = 300 units and *I* = 3 inputs, a time constant *τ* = 0.05*s*, and an integration time step *dt* = 0.01*s*. The noise *η* was randomly sampled from the Gaussian distribution *𝒩* (0, 0.2) for each time step. Each unit had an offset bias, 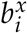, which was initially set to zero. The initial states *x*_*t*=0_ were sampled from the uniform random distribution *𝒰* (*−*0.2, 0.2). All networks were fully recurrently connected, with the recurrent weights *J*_*ij*_ initially sampled from the Gaussian distribution 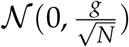, where *g* = 1.2. The time-dependent inputs *s* fed into the network had input weights *B* initially sampled from the uniform distribution *𝒰* (*−*0.1, 0.1). These inputs consisted of a one-dimensional fixation signal which started at 2 and went to 0 at the go cue, and a target signal that remained at 0 until the visual cue was presented. The two-dimensional target signal (2 *cos θ*^*target*^, 2 *sin θ*^*target*^) specified the reaching direction *θ*^*target*^ of the target.

The networks were trained to produce two-dimensional outputs *p* corresponding to *x* and *y* positions of the experimentally recorded reach trajectories, which were read-out via the linear mapping:

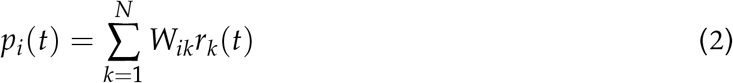

where the output weights *W* were sampled from the uniform distribution *U* (−0.1, 0.1).

#### Training

Networks were trained to generate positions of reach trajectories using the Adam optimiser^72^ with a learning rate *l* = 0.0005. The loss function *L* was defined as the mean squared error between the two-dimensional output and the target positions over each time step *t*. The first 50 time steps were not included to allow network dynamics to relax.

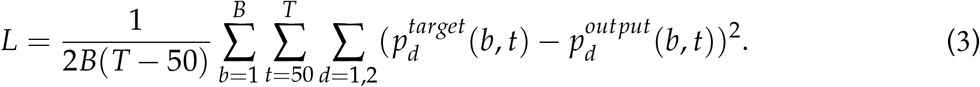

To examine whether two networks could have different latent dynamics while producing the same motor output, we devised a network with additional constraints to perform the behavioural task with distinct latent dynamics. We added a loss term that penalised the CC between the latent dynamics of the network being trained and those of another previously trained ‘standard’ network during movement execution:

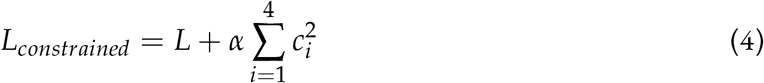

where *c*_*i*_ is the *i*-th CC and *α* = 0.5.

Networks were trained until the average loss of the last ten training trials fell below a threshold of 0.2cm^2^, for at least 50 and up to 500 training trials, with a batch size *B* = 64. Each batch had equal numbers of trials for each reach direction. Both standard and constrained training were performed on ten different networks initialised from different random seeds.

## Acknowledgements

We thank Kevin Mitchell for suggesting the concept of the genome encoding a ‘generative model’ and other helpful comments regarding this research, Leo Li for his contribution to the decoder analysis, and Carolina Massumoto for the monkey and mouse illustrations. This work was supported in part by: grant H2020-MSCA-IF-2020-101025630 from the Commission of the European Union (M.S.), grant 108908/Z/15/Z from the Wellcome Trust (J.C.), grants NS053603 and NS074044 from the NIH National Institute of Neurological Disorders and Stroke (L.E.M.), grant (chercheurs-boursiers en intelligence artificielle) from the Fonds de recherche du Québec Santé (M.G.P.), grant EP/T020970/1 from the UKRI Engineering and Physical Sciences Research Council (J.A.G.), and grant ERC-2020-StG-949660 from the European Research Council (J.A.G.).

## Author contributions

M.S., M.G.P., and J.A.G. devised the project. J.P. and J.T.D. provided the mouse datasets. M.G.P. and L.E.M. provided the monkey datasets. M.S. and J.C.C. analysed data and generated figures. M.S., J.C.C., J.T.D., M.G.P., J.A.G. interpreted the data. M.S., J.C.C., M.G.P., and J.A.G. wrote the manuscript. All authors discussed and edited the manuscript. M.G.P. and J.A.G. jointly supervised the work.

## Code availability

All analyses were implemented using custom python code using open-source software. All the result panels are reproducible by running Jupyter notebooks. Code to reproduce all the results is openly available in https://github.com/BeNeuroLab/2022-preserved-dynamics.

## Data availability

Many of the monkey datasets are publicly available on Dryad (https://datadryad.org/stash/dataset/doi:10.5061/dryad.xd2547dkt). The remaining data will be made available upon reasonable request.

## Supplementary Material

**Supplementary Figure S1:**
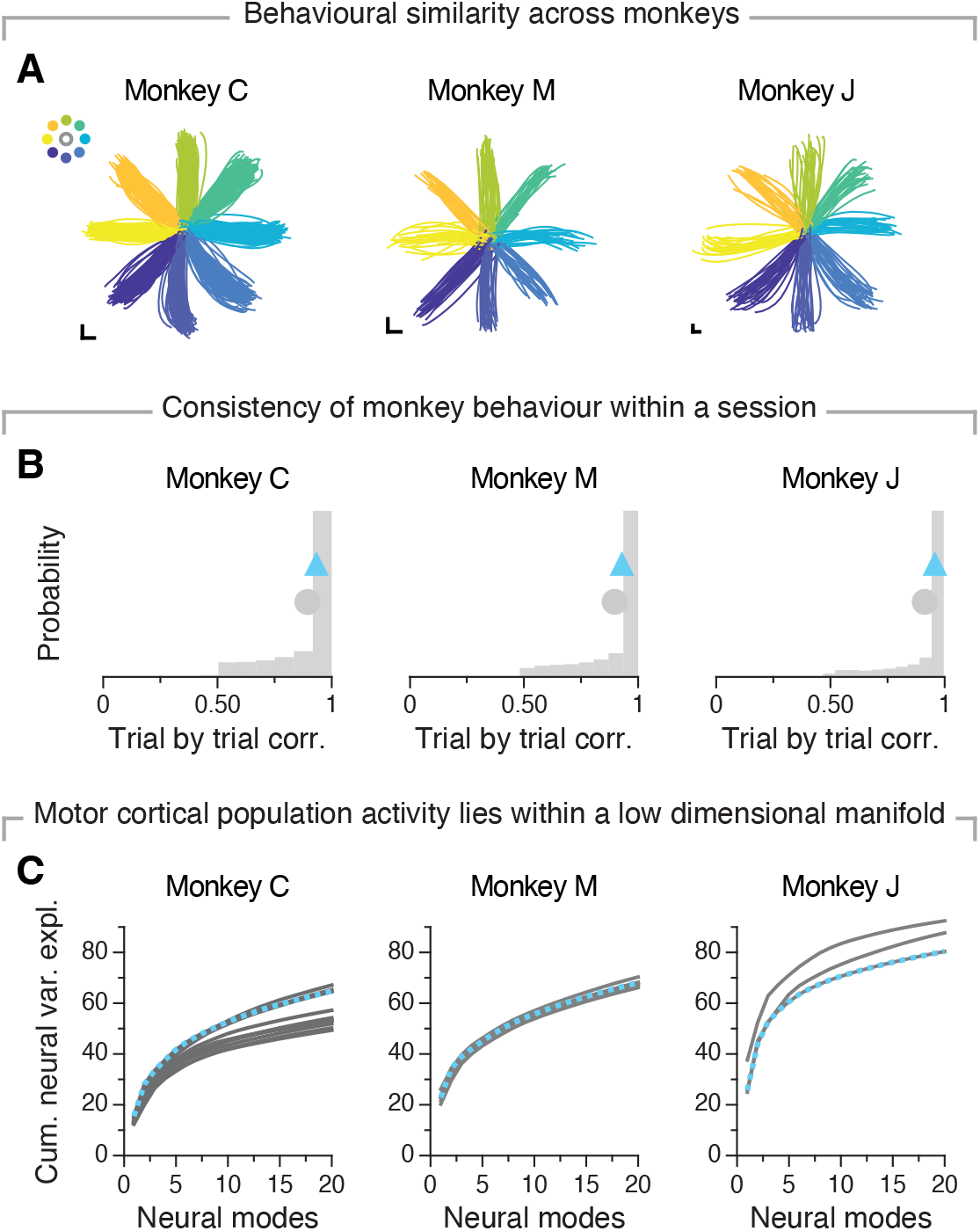
Behavioural and neural data from monkeys performing a centre-out reaching task. **A)** Hand trajectories of a single session of every monkey included in this work. Each line represents one trial, colour coded by target (inset). Scale bar, 1 cm. **B)** Distribution of the behavioural correlations for all pairs of trials from every included session for each of the 3 monkeys. Grey circle, mean. Blue triangle, mean for the representative session shown in Panel A. **C)** Cumulative neural variance explained as a function of the number of neural modes (principal components) included. Each line, one session. Dashed blue line, the same session shown in Panel A.

**Supplementary Figure S2:**
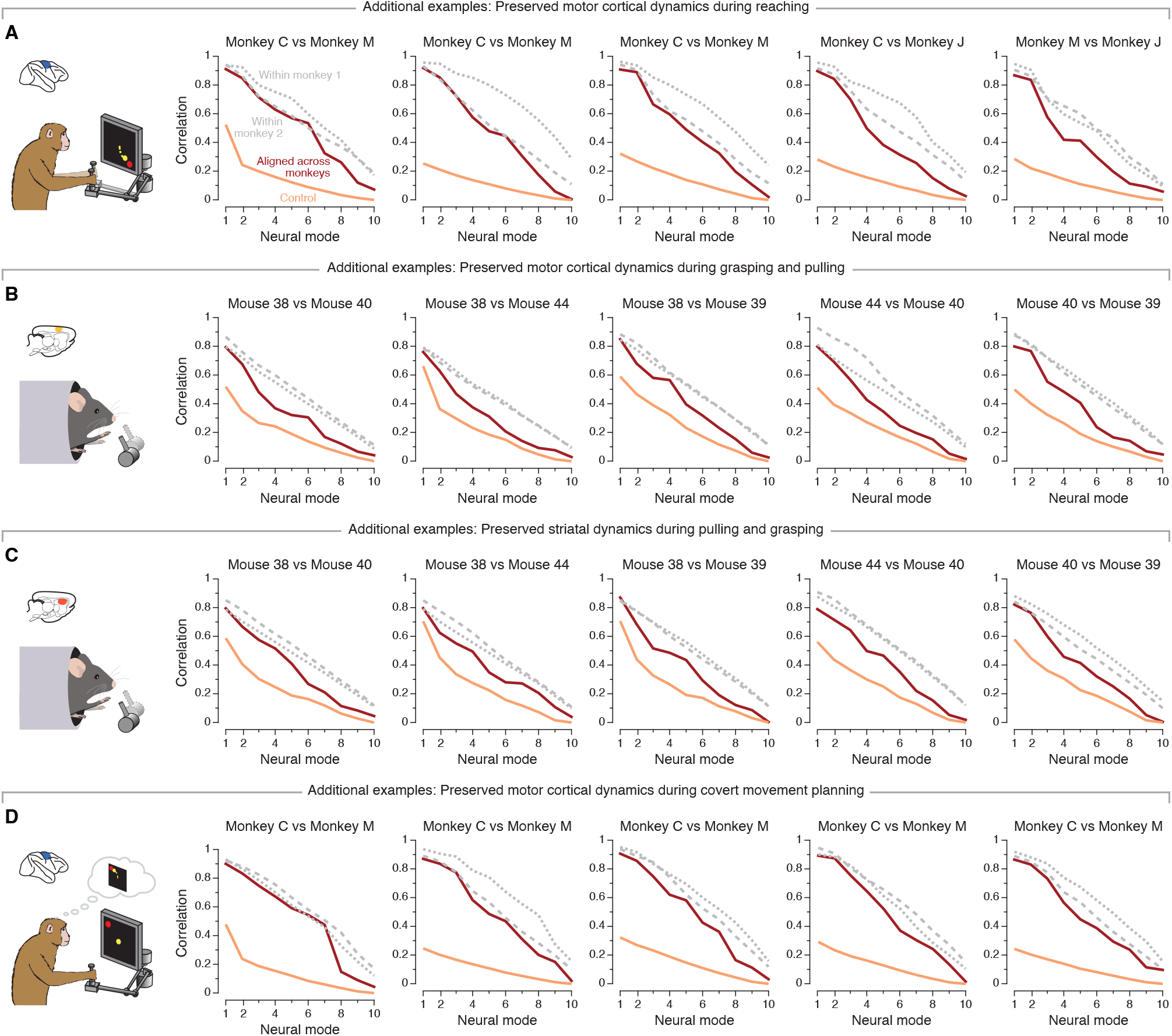
Example canonical correlations between pairs of sessions from two different monkeys and mice. **A)** Example canonical correlations between motor cortical latent dynamics for monkeys during movement execution. Shown are five of the individual comparisons included in the pooled results in Figure 2D. **B)** Example canonical correlations between motor cortical latent dynamics for mice during movement execution. Shown are five of the individual comparisons included in the pooled results in Figure 3B. **C)** Same as Panel B for mouse striatum. **D)** Same as Panel A but during movement planning.

**Supplementary Figure S3:**
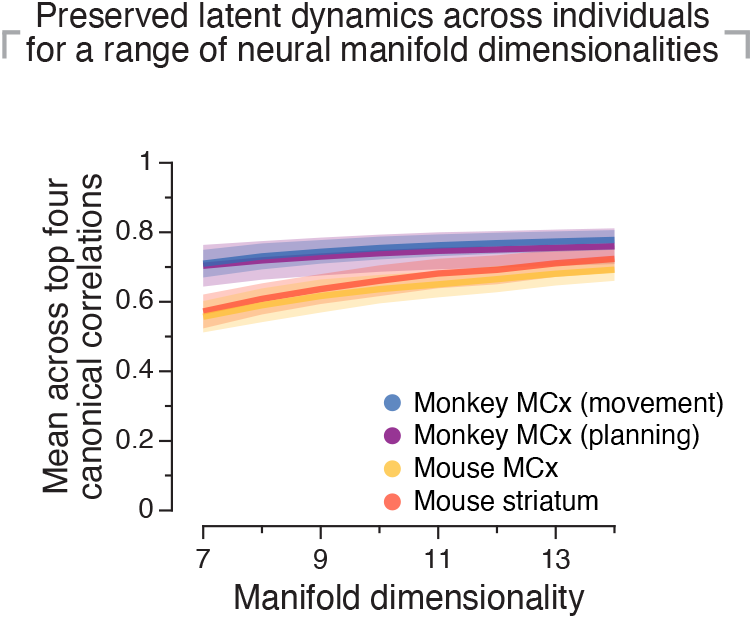
Preservation of latent dynamics does not depend on the number of manifold dimensions. Average canonical correlation as function of the number of manifold dimensions (principal components) for each of the four datasets considered (legend). Data pooled across all comparisons separately for each dataset. Line and shaded surface, mean*±*s.d. across the top four canonical correlations. MCx, motor cortex.

**Supplementary Figure S4:**
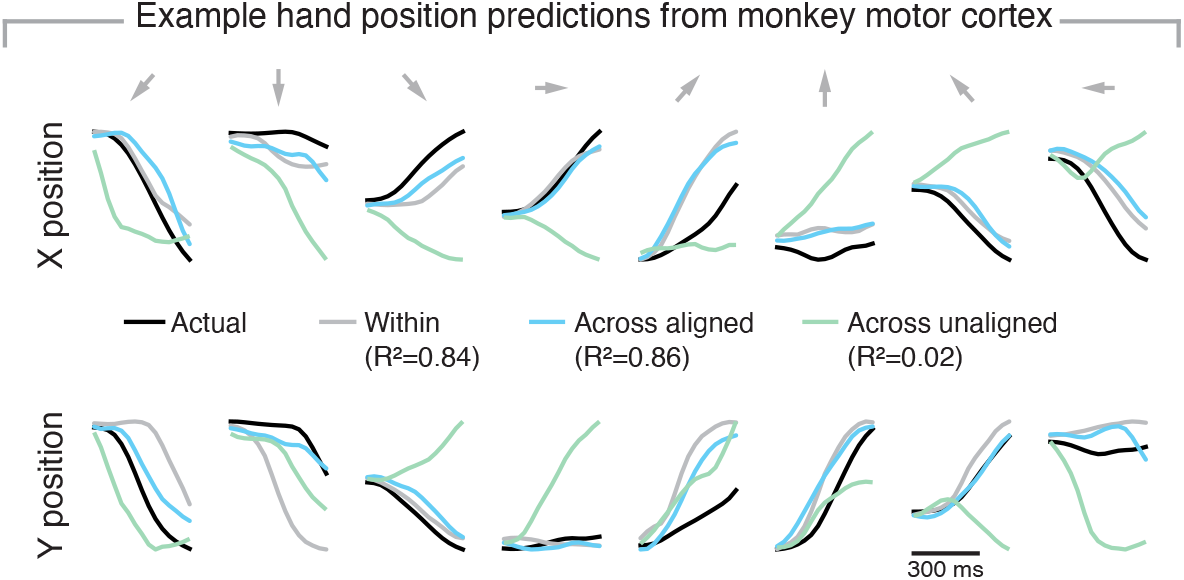
Example kinematic predictions illustrate that the preserved latent dynamics across animals are behaviourally relevant. Actual and decoded hand trajectory during one reach to each of the eight targets along the *X* (top) and *Y* directions (bottom); arrows on top of each column indicate target direction. The figure shows predictions using three models along with their respective accuracy (legend). Data from the same representative session shown in Figure 2B. Note that a decoder trained on a different monkey after alignment of their latent dynamics (‘Across aligned’) performed virtually as well as a decoder trained and tested on the same session from the monkey being tested (‘Within’).

**Supplementary Figure S5:**
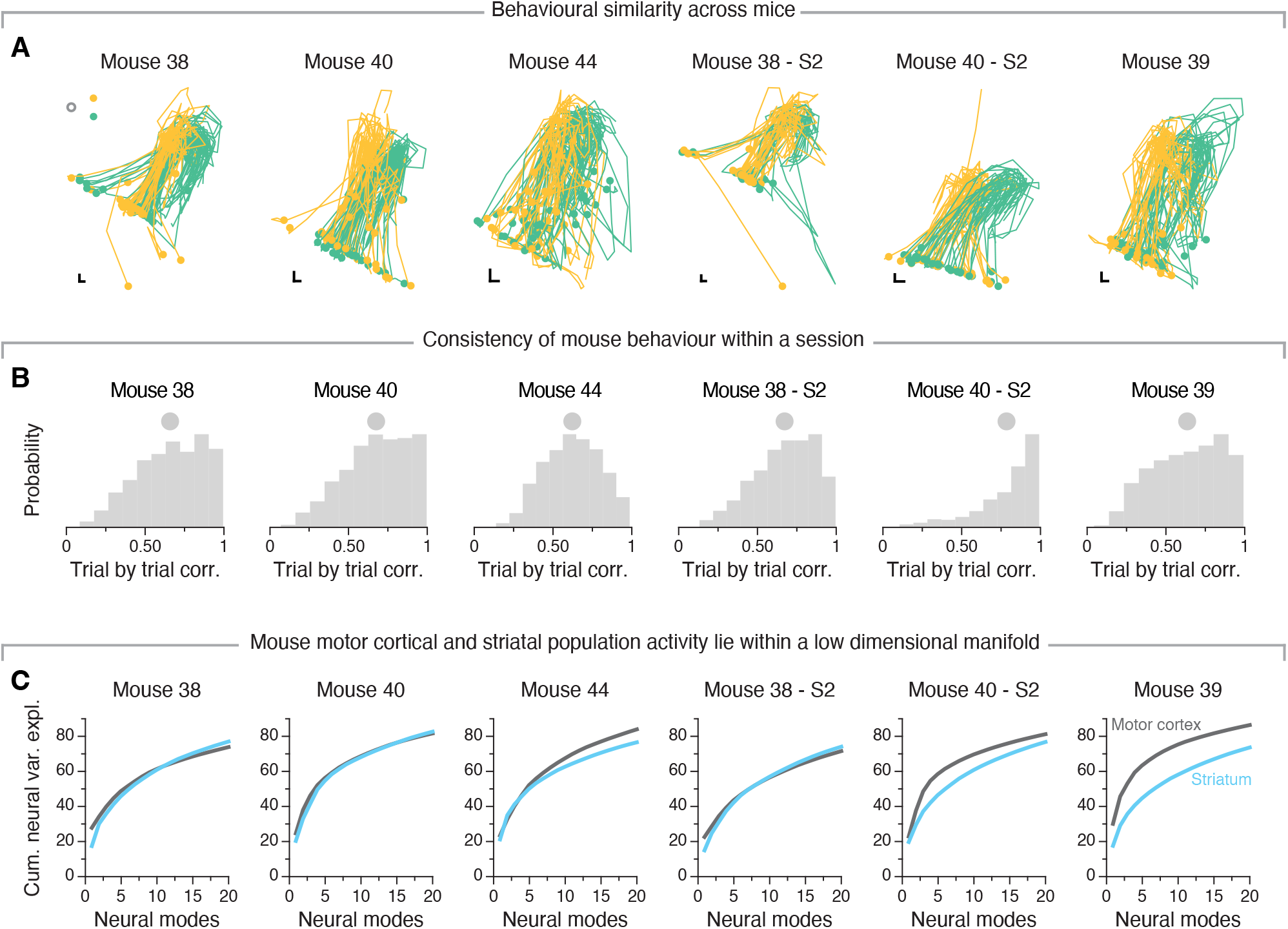
Behaviour and neural activity for mice performing a grasping and pulling task. **A)** Hand trajectories of every session included in this work (notice that two mice have two sessions each). Each line represents one trial, colour coded by target (left, yellow; green, right; see inset). Circle, initial hand position. Scale bar, 1 mm. **B)** Distribution of the behavioural correlations for all pairs of trials from every session. Grey circle, mean. **C)** Cumulative neural variance explained as a function of the number of neural modes (principal components) included, colour coded by brain region.

**Supplementary Figure S6:**
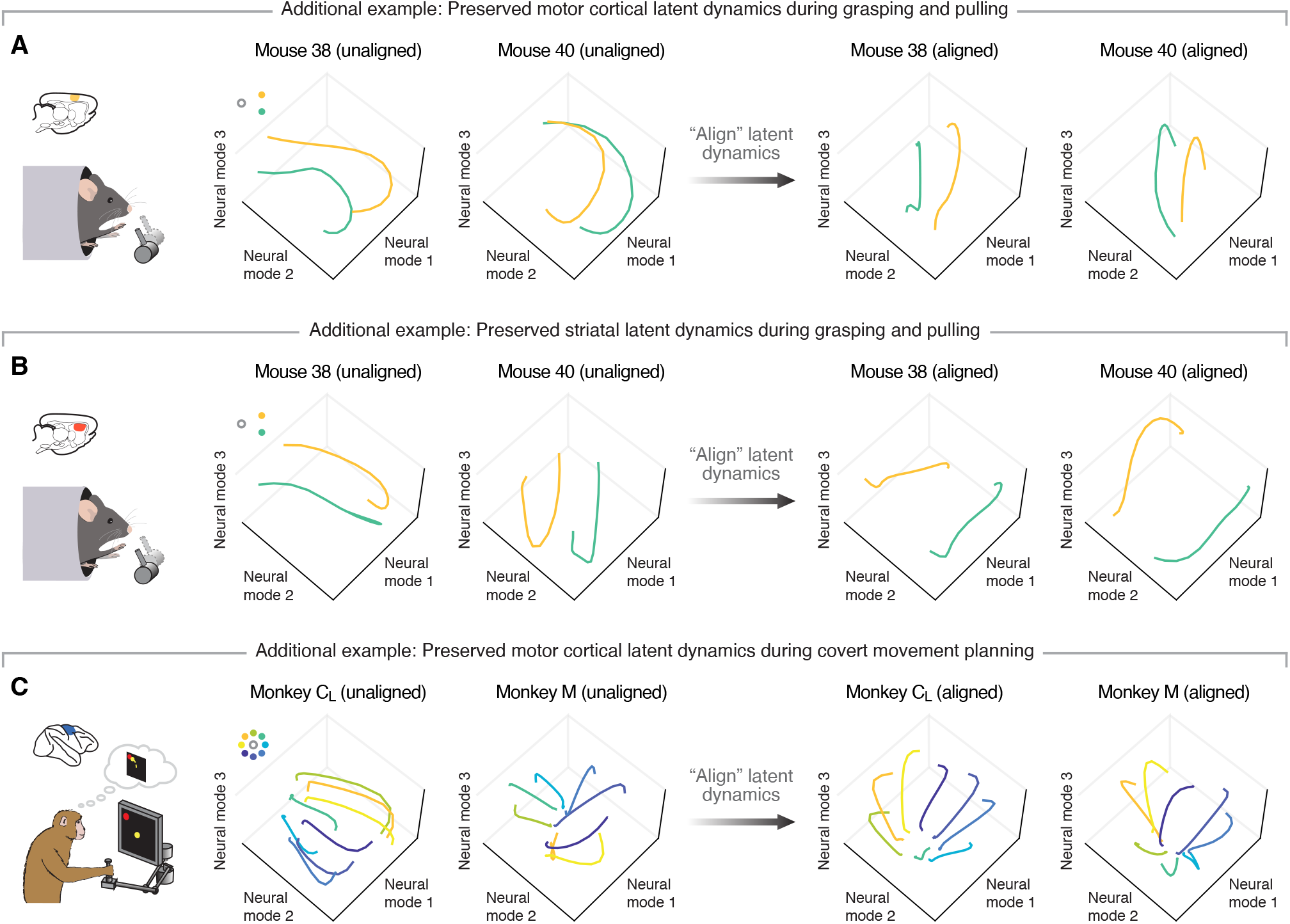
Additional Examples of preserved latent dynamics in monkeys and mice. **A)** Trajectories described by the motor cortical latent dynamics before (left) and after (right) alignment for a representative pair of sessions from two different mice. Individual lines, mean latent trajectory to each target (colour coded as in the inset). **B)** Similar to Panel B, for striatal latent dynamics in mice. **C)** Similar to Panel C, for motor cortical latent dynamics during covert movement preparation in monkeys.

**Supplementary Figure S7:**
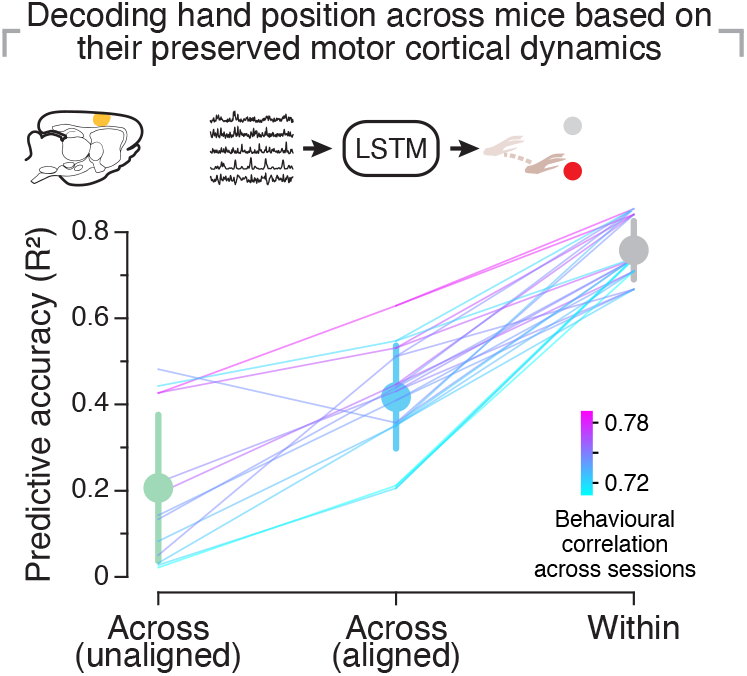
Across-mouse decoding of hand position based on the aligned latent dynamics. Decoders trained on the aligned latent dynamics from one mouse allowed predicting hand position from a different mouse with reasonable accuracy (blue), considerably better than decoders based on the unaligned latent dynamics (green). Lines, individual comparisons between two sessions from different mice (*n* = 6 sessions and *n* = 13 pairs), colour coded based on the correlation in hand kinematics across them (legend). Errorbars, mean*±*s.d.

**Supplementary Figure S8:**
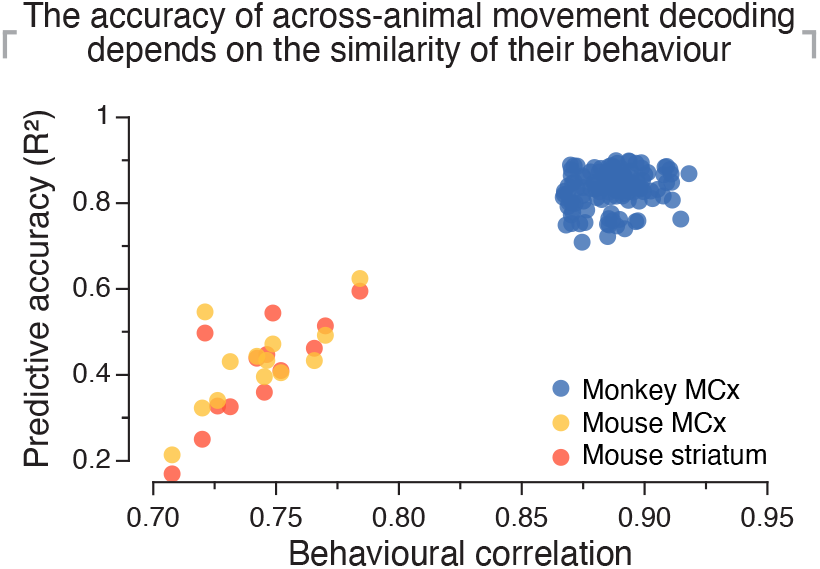
The accuracy of across-animal decoders is constrained by the similarity of their behaviour. Decoding performance based on the aligned latent dynamics for each pair of sessions for the three movement execution datasets (legend) as function of their mean behavioural correlation. Similar to Figure 2I.

**Supplementary Figure S9:**
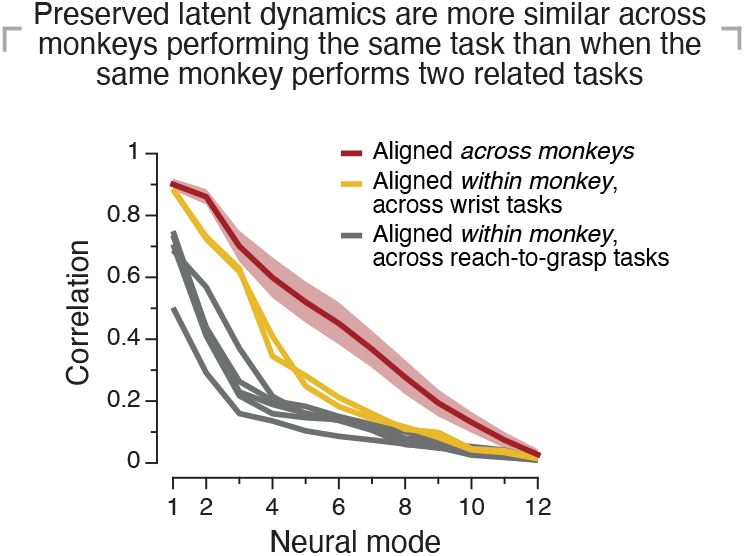
The latent dynamics of two different monkeys performing the same task are more correlated than those of the same monkey performing two related tasks. The across monkey results (red) illustrate preservation of the latent dynamics as monkeys performed the same centre-out reaching task. The results are the same as in Figure 2E only regenerated with 12 components to match the across-task analysis in Ref.^27^. The across-task comparison illustrates the similarity of the latent dynamics generated as monkeys performed two distinct but related wrist tasks (including isometric force generation, movement without a load, and elastic-loaded movement), or two distinct but related reaching and grasping tasks (a power grasp task, and grasping, transporting and dropping a ball); adapted from Supplementary Figure 7 in Gallego *et al*.^27^. *Each yellow and dark grey trace, one session*.

**Supplementary Figure S10:**
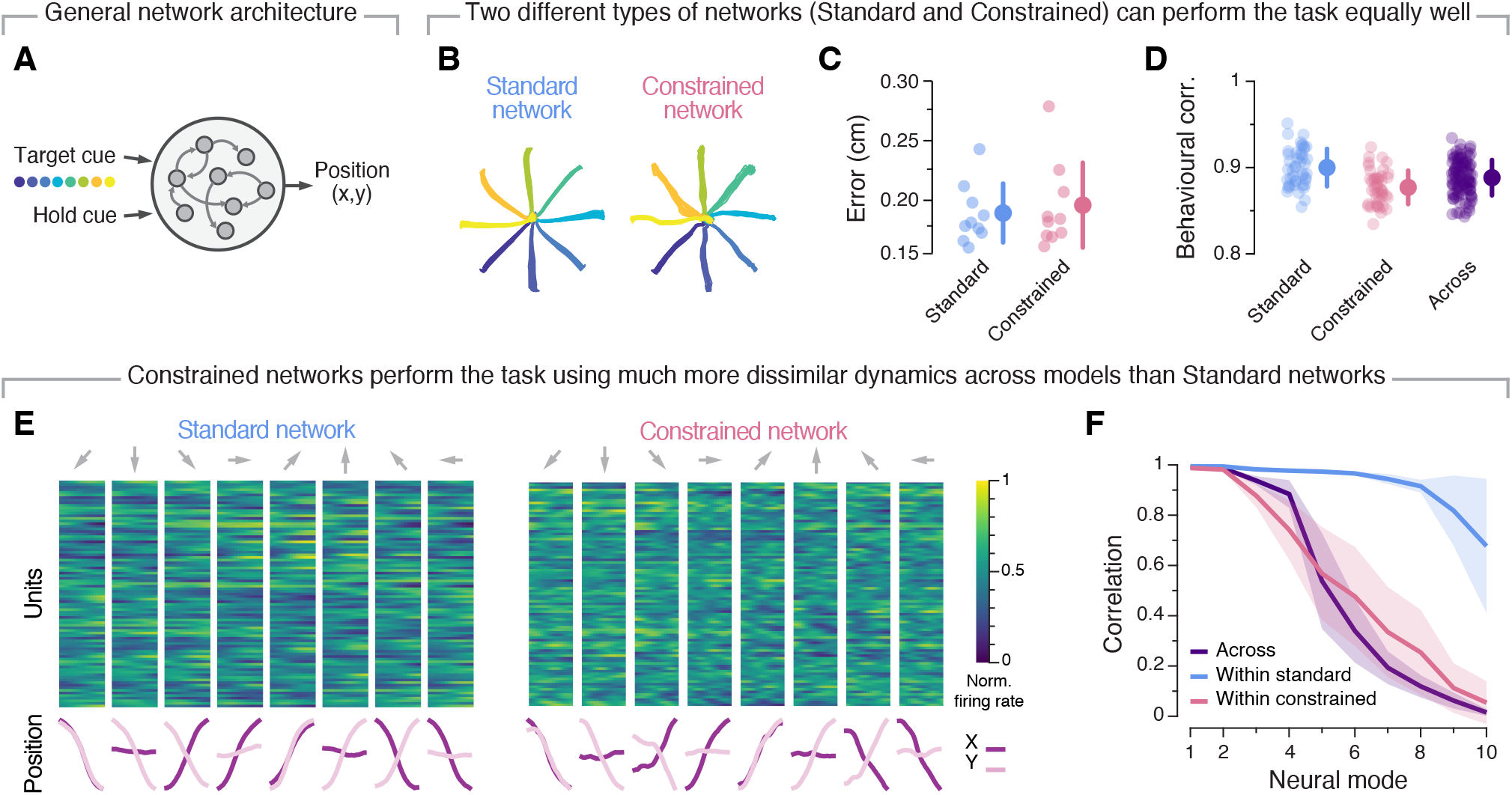
Behavioural similarity is necessary but not sufficient for the preservation of latent dynamics: different types of recurrent neural networks (RNN) that perform the same reaching task equally well can generate dissimilar latent dynamics. **A)** RNNs were trained to perform the centre-out reaching task we studied in monkeys. **B)** Output of example networks for the two groups of RNNs we trained: ‘Standard’ networks (left), and ‘Constrained’ networks that were penalised for canonical correlation similarity to the latent dynamics of Standard networks (right). **C)** Mean-squared error between network output and target positions following training for each type of network. **D)** Behavioural correlation across pairs of Standard and Constrained networks, and across networks from these two different types. Markers, individual comparisons. Errorbars, mean ± s.d. **E)** Generated network dynamics and behaviour for each reach direction for example Standard (left) and Constrained (right) networks. Data presented as in Figure 2B. **F)** Similarity in latent dynamics between pairs of Standard networks (blue) and Constrained networks (pink), and between pairs of networks for each type (purple). Note that Constrained networks have correlations that are much more dissimilar than Standard networks, suggesting that behavioural similarity is not sufficient to obtain high correlations.

